# PSEN1 mutant marmoset fibroblasts mimic multi-omic signatures of Alzheimer’s disease

**DOI:** 10.64898/2026.04.24.720222

**Authors:** Sonal Kumar, Annat Haber, Catrina Spruce, Duc Duong, Nicholas T. Seyfried, Lauren Bailey, Sang-Ho Choi, Stephanie Hachem, Yongshan Mou, Seung-Kwon Ha, Jung Eun Park, Gregg E. Homanics, Stacey J. Sukoff Rizzo, Afonso C. Silva, Gregory W. Carter

**Author notes:** **Correspondence should be directed to:** Gregory W. Carter, PhD The Jackson Laboratory, 600 Main Street, Bar Harbor, ME 04609, USA.

## Abstract

**INTRODUCTION:** The slow, age-related development of Alzheimer’s disease (AD) and inaccessibility of early-stage brain tissue necessitates model studies to understand its origins and progression. Non-human primate models can provide a platform for linking molecular changes to translatable phenotypes. Here, we assess fibroblast lines derived from marmosets with engineered variants in the *PSEN1* gene.

**METHODS:** Fibroblast cultures were obtained from 10 animals and assayed using a NanoString AD gene expression panel and label-free proteomics. We compared mutant expression changes to human AD signatures in human iPSC-derived neurons and postmortem brains to assess disease relevance.

**RESULTS:** Gene products involved in amyloid-beta interaction and regulation were differentially expressed, providing evidence for the functional relevance of the engineered fibroblasts. Both gene and protein expression changes in the undifferentiated fibroblasts correlated with human iPSCs from AD donors reprogrammed into neuronal lineages, as well as postmortem brains derived from case-control cohorts. Altered expression profiles were noted based on marmoset donor sex and mutation status, highlighting underlying sex-specific biology relevant to Alzheimer’s disease.

**DISCUSSION:** These findings demonstrate that disease-relevant pathways and processes are altered in fibroblasts from mutant marmosets, emphasize the complementarity of transcriptomic and proteomic profiling in AD, and provide a roadmap for more advanced molecular studies of AD in aging marmosets and marmoset-derived cell models.

## 1. INTRODUCTION/BACKGROUND

The proliferation of genetic and multi-omic studies of Alzheimer’s disease and related dementias (ADRD) has generated abundant molecular hypotheses of disease origins and progression. While analogous studies in mouse models have proven useful in understanding the biology that underlies these molecular changes^1–4^, the absence of spontaneous AD-related neuropathology, lack of frank neurodegeneration in response to amyloid deposition, and restricted behavioral repertoire have limited translational relevance. This motivates the establishment of a non-human primate model that spontaneously develops age-dependent AD neuropathologies and exhibits a more complete form of early-onset dementia when carrying familial AD genetics. The common marmoset *Callithrix jacchus*, a small new world monkey with an experimentally tractable lifespan, potentially balances translatability and practicality in a laboratory model^5,6^. We recently introduced mutations in the *PSEN1* gene in marmosets using CRISPR/Cas9 genetic engineering^7^. Elevated amyloid levels were observed in plasma of mutation carriers versus wild-type controls^7^. To assess molecular changes arising from these perturbations, here we assess gene and protein expression in cells derived from these animals and unmanipulated, age-matched controls.

The use of transcriptomic, proteomic, and other omics-scale molecular assays in human and animal studies enables multi-dimensional alignment between human disease and experimental systems that seek to model aspects of ADRD. This deep characterization has identified the specific disease-relevant changes in mouse^3,4,8,9^ and cell models^10^, specifying the aspects of the disease best captured by each model. Pairing human cohort studies with induced pluripotent stem cell (iPSC) lines derived from study individuals further enables modeling of how distinct cell types may contribute to disease at various stages of initiation and progression. Following this approach, here we assess fibroblast cultures derived from marmosets skin biopsies and compare their multi-omics profiles with human-derived cell lines differentiated into neurons from the ROSMAP-IN study^11^. We demonstrate changes in both transcript and protein levels that are conserved in the mutant animals and neuronal cells derived from induced pluripotent stem cells (iPSCs) taken from human AD subjects and age-matched controls. We further compared transcriptomics and proteomics signatures to postmortem human brain studies of late-onset AD^12,13^ and identified a subset of disease alterations recapitulated in the marmoset fibroblasts. These conserved changes are present despite the lack of differentiation of the fibroblasts to similar neuronal lineages, suggesting fundamental alterations that are driven solely by *PSEN1* mutations. This analysis provides a basis for future multi-omics studies of marmoset tissues and marmoset-derived cell models.

## 2. METHODS

### 2.1 Marmoset fibroblast samples

All animal experimental procedures were conducted under state and federal laws, locally approved by The University of Pittsburgh Institutional Animal Care and Use Committee (IACUC protocol 24014391), and were in line with and strictly adhered to the Guide for the Care and Use of Laboratory Animals. Subjects were socially housed in a dedicated marmoset AAALAC-accredited facility at the University of Pittsburgh and maintained at a temperature range of 76–78 °F and 30–70% humidity on a twelve-hour light/dark cycle. Subjects were fed a diet consisting of twice daily provisions of commercial chow (Callitrichid High Fiber Diet #5LK7, Lab Diet; Marmoset Diet TD#130059, Envigo Teklad, Madison WI, USA), supplemented with various fruit and vegetable treats and health monitored daily by the veterinarian staff and veterinarians. Marmoset fibroblasts were obtained by skin biopsy cultures from five *PSEN1* mutant marmosets and age- and sex-matched non-carrier control marmosets, for a total of ten animals (**Table 1**). Skin fibroblasts were used to avoid potential chimerism in cells and tissues of hematopoietic origin as a consequence of genetic exchange occurring between siblings in utero^14^. *PSEN1* variants C410Y and A426P were targeted by CRISPR/Cas9 editing, resulting in a variety of *PSEN1* genotypes in the founder animals, the full details of which were previously described^7^, with additional data available on the Alzheimer’s Disease Knowledge Portal^15^. Animals were anesthetized for obtaining skin biopsies, using 10 mg/kg ketamine IM, followed by 2% isoflurane in 100% O_2_ inhalation for the remainder of the procedure. Small pieces of skin tissues were washed and minced in Hank’s Balanced Salt Solution (HBSS, Gibco) and cultured in Iscove’s Modified Dulbecco’s Medium (IMDM, Gibco) containing 10% fetal bovine serum (FBS, Gibco), 1% penicillin/streptomycin (Gibco), and 1mM Glutamax (Gibco) at 38 °C, in a humidified atmosphere of 5% CO_2_ and 95% air. A fibroblast monolayer from the tissue explants was established after 10-14 days. At passage zero, the cells were cryopreserved in Bambanker freezing media (GC Lymphotec Inc., Tokyo, Japan). The media was changed every 2-3 days during the culture period until they were confluent and the cells from passage numbers 1 to 4 were retrieved from the monolayer by trypsinization.

**Table 1:** Individual specifications for the marmoset samples included in this study.

| Individual ID | Sex | Sampling age<br>(months) | Targeted mutation and genotype |
| --- | --- | --- | --- |
| 4 | Male | 8.2 | PSEN1-C410Y_Y410/+1 |
| 5 | Male | 8.2 | PSEN1-C410Y_Y410/Y410 |
| 1 | Male | 6.7 | PSEN1-A426P_P426/D20 |
| 6 | Female | 8.4 | PSEN1-C410Y_Other |
| 105 | Female | 2.8 | PSEN1-A426P_P426/Indel |
| 101 | Female | 3.5 | Wild-type |
| 9 | Male | 8.2 | Wild-type |
| 2 | Male | 7.2 | Wild-type |
| 3 | Male | 8.2 | Wild-type |
| 8 | Female | 7.2 | Wild-type |

Metadata files for individual animals (https://www.synapse.org/Synapse:syn63926850; syn63926850) and biospecimen samples (https://www.synapse.org/Synapse:syn63927118; syn63927118) are available on the AD Knowledge Portal^15^.

### 2.2 NanoString assay and data preprocessing

The NanoString Human AD gene expression panel was used for gene expression profiling on the nCounter platform (NanoString, Seattle, WA, USA), as described previously^8^. This panel was originally designed to maximize coverage of Alzheimer’s disease gene modules derived from postmortem brains in three cohorts^12^. NanoString nSolver software was used to generate gene expression counts. Normalization with positive probes was carried out for endogenous gene expression, by using sample-specific normalization factors (mean-centered probe expression normalized to each sample). The gene counts were then log_2_ transformed for further analysis (https://www.synapse.org/Synapse:syn66641748; syn66641748).

### 2.3 Label free mass spectrometry proteomics and data preprocessing

Samples were prepared using the EasyPep MS Sample Preparation Kit (Thermo Fisher Scientific, #A40006) according to the manufacturer’s protocol. Briefly, samples were resuspended in 175 μL of lysis buffer, followed by the addition of 2 μL of nuclease. To aid in lysis, samples were aspirated 10 times and then spun down. Protein concentration was determined using a BCA assay, and the equivalent of 100 μg protein from each sample was aliquoted for downstream processing. Samples were then normalized to a final volume of 100 μL with lysis buffer, followed by reduction and alkylation at 95 °C for 10 minutes. A LysC and trypsin enzyme mix was added according to protocol, and digestion was carried out for 3 hours. Samples were then cleaned up and eluted with 300 μL of elution buffer. A 20 μL aliquot was taken and dried separately. All eluants were dried down to completion via a speed vacuum concentrator (LabConco).

Dried peptides (20 μL eluant aliquots) were resuspended in 15 μL of loading buffer (0.1% formic acid) and 4 μL (approximately 2 μg) of sample was separated using a Dionex RSLCnano liquid chromatography (LC) system (Thermo Fisher Scientific) fitted with a self-made 25-cm-long, 150 μm internal diameter fused silica column packed with 1.7 μm Water’s CSH resin. The LC flowrate was set at 1.5 μL per minute and went from 7 to 40% buffer B (buffer A: Water with 0.1% formic acid and buffer B: Acetonitrile with 0.1% formic acid). The gradient started at 7% to 40% B in 32 minutes, and was followed by a three-minute flush with 99% B. The mass spectra were collected on an Orbitrap Eclipse Tribrid mass spectrometer equipped with a FAIMS Pro ion mobility interface (Thermo Fisher Scientific). Each mass spectrometer cycle consisted of two compensation voltages (−40 and −60) each, with one full survey scan followed by as many tandem mass spectrometer scans that could fit within one second. The full survey scan was collected in the orbitrap at a resolution of 120,000 with standard automatic gain control (AGC) and automatic injection timings. High energy collision dissociation (HCD) tandem scans were collected in the ion trap with an isolation width of 1.6 mass to charge, collision energy of 35%, and scan rate set to rapid. Dynamic exclusion was set to exclude ions for 60 seconds once they had been selected for fragmentation.

Proteomics raw files were analyzed using Proteome Discoverer (v2.4, Thermo Fisher Scientific). All spectra were searched against a 2020 *Callithrix jacchus* database consisting of 48,340 reference protein sequences. Search parameters included fully tryptic digestion, maximum miscleavages of 2, a precursor ion mass tolerance of 10 ppm, fragment ion tolerance of 0.6 Daltons, dynamic modification of oxidized methionine (+15.995 Da), and static modification of carbamidomethyl cysteine (+57.021). All peptide spectral matches were filtered through Percolator^16^ and for label free quantification (LFQ), peak areas were matched and calculated using the Minora feature detection node.

Normalized protein abundances were log_2_ transformed for subsequent analyses (https://www.synapse.org/Synapse:syn64544741; syn64544741). Annotations obtained from UniProt.org (downloaded in August 2025) were used to map protein accession IDs to protein names for cross-species comparison. The Retrieve/ID Mapping functionality was used for given accession IDs, and a customized output containing the columns protein names and primary gene names was downloaded for our use. 2172 unique peptide IDs were reduced to 2123 unique proteins after removing duplicated gene symbols.

### 2.4 ROSMAP-IN RNA-Seq and proteomics data and preprocessing

All transcriptomic and proteomic data, as well as metadata for the human samples were obtained from a study of differentiated neurons from induced pluripotent stem cells (iPSCs) derived from human donors in the Religious Orders Study and Memory and Aging Project (ROSMAP)^11^. Metadata from the ROSMAP iPSC-Derived Induced Neurons (ROSMAP-IN) study was obtained from the AD Knowledge Portal (https://www.synapse.org/Synapse:syn25954752; syn25954752; https://www.synapse.org/Synapse:syn25958845; syn25958845). Processed ROSMAP-IN RNA-Seq data in transcripts per million (TPM) was obtained from the AD Knowledge Portal (https://www.synapse.org/Synapse:syn25954748; syn25954748). TPM counts were then log_2_ transformed, and genes with log_2_ TPM below three in all samples were filtered out as below quantifiable expression. Duplicated gene entries were removed. Label-free LC-MS/MS proteomics were obtained as log_2_ transformed values with filtering done to remove proteins with seven or more missing values for a total of 40 samples (https://www.synapse.org/Synapse:syn25954751; syn25954751). Protein accession IDs were replaced with primary gene names as described in Section 2.3 (Accessed May 2025). Full assay methods and results were previously published^11^.

### 2.5 Differential expression analysis

For both omics modalities in each model system, we determined the effect of each experimental factor by fitting a multiple regression model of the following form using the lm() function in base R for the general equation:

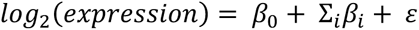

The factors for the marmoset datasets included sex and *PSEN1* genotype. Sex was encoded as a binary variable representing effects specific to males. *PSEN1* genotype was encoded with three levels 0 (wild-type), 1 (heterozygous), and 2 (homozygous for the target allele). For data from the ROSMAP-IN study, the factors sex, AD diagnosis, and *APOE2*, *APOE4* genotypes were encoded as binary variables indicating male, positive diagnoses of AD, and *APOE* genotype effects, respectively. CERAD and Braak score variables were scaled to values between 0 and 1. Differentially expressed genes in both cases were determined at a threshold of FDR < 0.1 using a Benjamini-Hochberg adjusted *p*-value for false discovery rate. The full model results for both marmoset NanoString and proteomics expression are given in **Supplementary Tables S1** and **S2**, respectively.

### 2.6 Comparison of ROSMAP-IN and marmoset omics

To compare gene expression and protein abundance effects, we computed correlations between regression coefficients (β) specific to each factor for both the human iNs and marmoset fibroblast cell lines across common genes and proteins. A subset of nominally significant genes (*p*-value < 0.05) for each ROSMAP-IN factor was chosen for comparing expression effects with corresponding marmoset genes. Correlations were calculated for this subset between each pair of human-marmoset factors to determine similarity of effects. Pearson’s correlation was implemented using the `cor.test()` function in base R, with statistical significance noted for correlations with *p* < 0.05.

### 2.7 Alignment to human AMP-AD co-expression RNA and protein modules

Data from the Accelerating Medicines Partnership for Alzheimer’s Disease (AMP-AD) RNA co-expression modules was obtained from the AD Knowledge Portal (https://doi.org/10.7303/syn11932957.1; syn11932957). To assess disease-specific gene expression patterns, we used 30 human brain co-expression modules from meta-analysis of postmortem brain transcriptomes from three AMP-AD cohorts, which were further classified into five consensus clusters based on broader cell-type specificity and functional annotations^12^. Viewing these modules as a summary of human transcriptomic changes in AD, we compared the effects of *PSEN1* mutations in marmoset NanoString data to human changes in AD cases versus controls. We note the human AD panel was designed to maximize coverage across these modules^8^. Correlations were calculated for gene effects per factor for the marmoset fibroblasts (β) and each module (residual log_2_ fold changes in AD cases versus controls after corrections for sex (reflecting male-specific effects), age, and technical covariates) using the `cor.test()` function in base R. Significant correlations were defined based on *p* < 0.05. We additionally compared proteomic changes across 44 co-expression modules built from tandem mass tag mass spectrometry (TMT-MS) data from the human dorsolateral prefrontal cortex (DLPFC), which were further annotated using gene ontology (GO) based analyses to identify the central biological theme for each module^13^. As with transcriptomic co-expression modules, correlations were calculated as previously described for the protein set common to those assessed for marmoset fibroblasts (β) and those in the protein modules (residual log_2_ fold changes in AD cases versus controls after corrections for sex, age, and technical covariates). Significant correlations were defined as *p* < 0.05.

### 2.8 Statistical analysis, data, and code availability

All tests were conducted in RStudio by Posit (2026.1.0.392) running R v4.5.0. More information on the analysis and experimental design is detailed above, and the code needed for reproducing the results and figures can be found on GitHub (https://github.com/sonalkumarr/marmoset_multiomics_pilot). Data hosted on the AD Knowledge Portal can be accessed after obtaining the necessary data use permissions through their unique synIDs.

## 3. RESULTS

### 3.1 *PSEN1* marmoset fibroblasts show similar trends for correlations between transcriptomic and proteomic changes as human iNs derived from AD cases

We used fibroblasts cultured from skin biopsies obtained from age and sex-matched *PSEN1* mutant and wild-type marmosets (n = 10) to assess molecular changes resulting from the engineered mutations. We measured the abundance of 770 genes using the NanoString Human AD gene panel and 2123 proteins as assessed by label-free quantification (LFQ) of liquid chromatography coupled tandem mass spectrometry (LC-MS/MS).

We first assessed the similarities in gene expression and protein abundance changes resulting from the *PSEN1* editing. This was compared to similar changes seen with Alzheimer’s disease in induced neuronal cell lines (iNs) derived from 53 human donors in the Religious Orders Study and Memory Aging Project (ROSMAP) cohorts. Multi-omic signatures from these iNs have identified proteins that show strong associations with corresponding pathology in donor postmortem brains including β-amyloid deposition, tau tangles, alterations in biological pathways, and measured cognitive decline.^11^

*PSEN1* effects (regression coefficients β; Methods) on gene and protein expression were correlated similarly in cell lines from both species (**Fig. 1**). For 1068 genes common to both RNA-seq and proteomics datasets for AD status in the ROSMAP iN samples, the Pearson’s correlation coefficient was 0.29 (*p* = 1.55 × 10^−21^). The more limited NanoString panel yielded 172 genes with corresponding protein abundances, for which the correlation was significant and similar to the human iNs (ρ = 0.27, *p* = 3.8 × 10^−4^). Only 60 genes were found common to both gene expression data sets.

**Figure 1:**
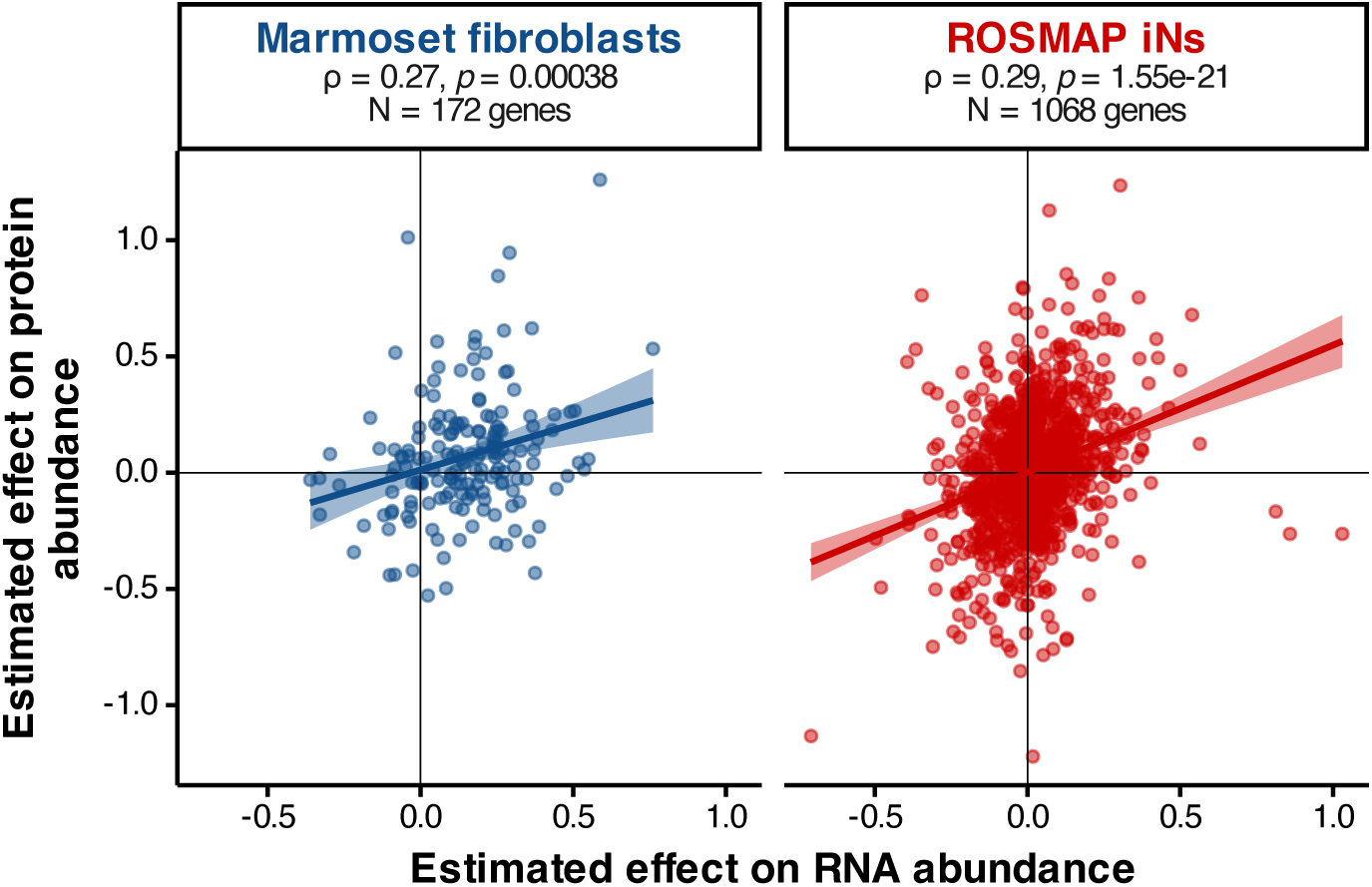
Alzheimer’s disease-related changes are conserved in transcriptomes and proteomes across human induced neuronal cell lines and marmoset fibroblasts. Correlations (ρ) between expression effects for genes (x-axis) and proteins (y-axis) for *PSEN1* genotype in marmoset fibroblasts (left) and AD diagnosis in ROSMAP induced neuronal cell lines (right). [ρ = Correlations computed using Pearson’s correlation coefficient; N = number of overlapping genes or proteins captured across the two omics modalities.]

### 3.2 Transcriptomic and proteomic signatures are significantly correlated between *PSEN1* genotype in marmoset fibroblasts and AD diagnosis in human iNs

We next compared differential gene and protein expression between both species *in vitro*, using the effects of the *PSEN1* mutations in marmoset fibroblasts and case-control diagnosis differences in human iNs. To determine the extent to which the marmoset *PSEN1* fibroblasts recapitulated significant effects in human iNs, we focused on genes and proteins that were differentially expressed for AD diagnosis in the human iNs (*p* < 0.05). Taking the subset of genes (N = 32) and proteins (N = 45) common to both species, we computed Pearson correlation coefficients between the effects of the *PSEN1* mutation in marmosets with the effects of AD diagnosis in iN donors. We found similar correlations that were statistically significant for the common genes (ρ = 0.35, *p* = 0.047) and proteins (ρ = 0.37, *p* = 0.012) (**Fig. 2A**). We note that these genes and proteins sets do not include any gene products in common, and the correlations are therefore driven by different gene/protein entities. These outcomes demonstrate distinct and translatable patterns for gene and protein expression in the *PSEN1* mutant cell cultures, even without differentiation to brain-resident cell types.

**Figure 2:**
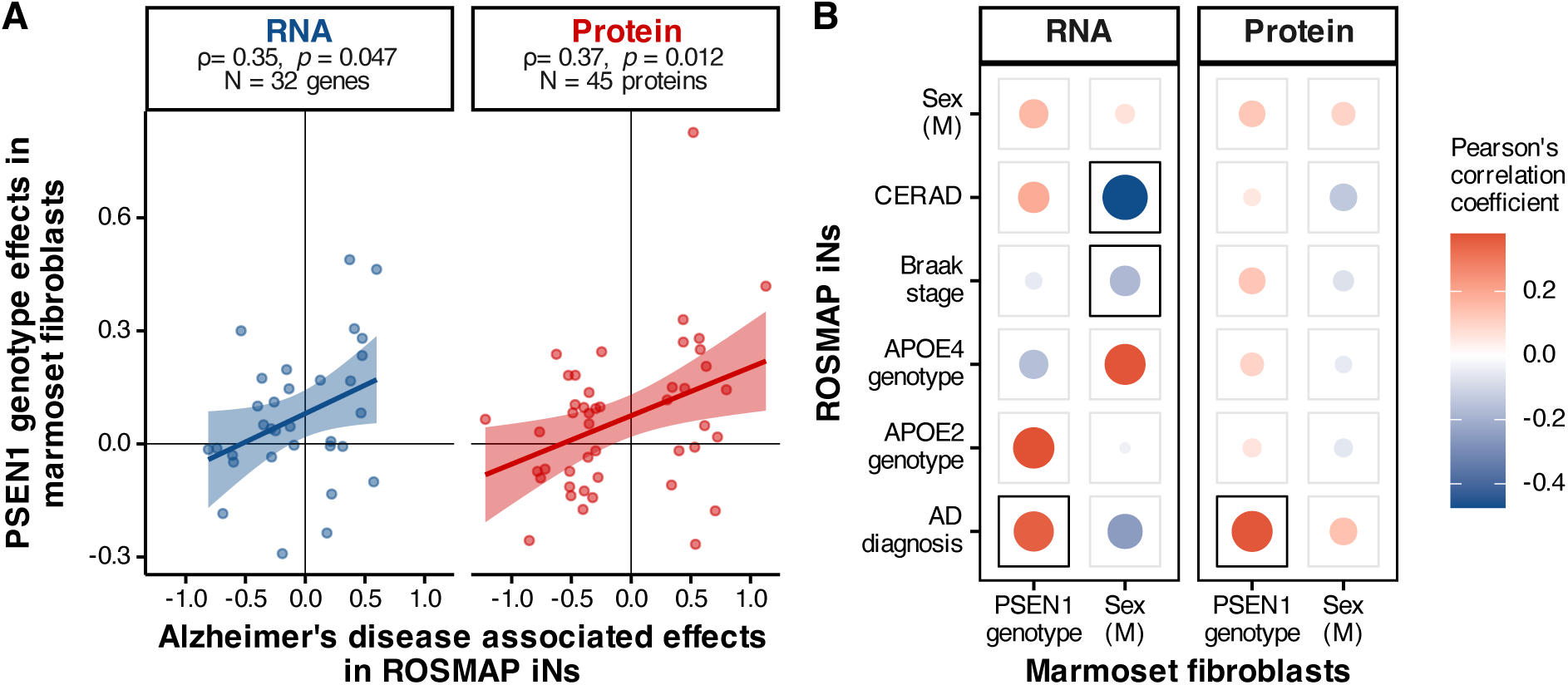
Human iNs from positive AD diagnoses and *PSEN1* marmoset fibroblasts have correlated gene and protein expression changes. **A.** Correlation between expression effects for AD status in human iNs (x-axis) and *PSEN1* genotype in marmoset fibroblasts (y-axis) for both transcriptomics and proteomics data. ρ denotes Pearson’s correlation coefficient. **B.** Correlations between expression effects calculated for all model factors for the marmoset fibroblasts (columns) and human iNs (rows). Disc size and intensity indicate the strength of the correlation, and significant (*p* < 0.05) correlations are framed.

While none of the 32 specified genes (**Fig. 2A**) were significantly differentially expressed (FDR < 0.1) in *PSEN1* mutant marmoset fibroblasts, two proteins out of the 45, namely PRKCSH (β = −0.14, FDR = 0.018) and SGTA (β = 0.25, FDR = 0.085), were downregulated and upregulated, respectively. PRKCSH, the beta subunit of glucosidase II, is involved in N-linked glycosylation in the endoplasmic reticulum^17^. This protein has been shown to colocalize with β-amyloid plaques and neurofibrillary tangles of tau in the hippocampus, with decreased protein levels observed in the AD brain^18^, with another study suggesting that PRKCSH may potentially be driving excitotoxicity mediated atrophy in the hippocampus and amygdala in AD^19^. SGTA, or the small glutamine rich tetratricopeptide repeat co-chaperone alpha protein, is integral to many biological processes including cell cycle, apoptosis, and synaptic transmission^20^. Direct evidence explaining SGTA’s role in Alzheimer’s disease is sparse; however, it has been shown to colocalize with intracellular aggregates in neurodegenerative diseases like Huntington’s^21^ and significant alterations in protein levels have been noted in other dementias with proteinopathic aggregates like Lewy-body and Parkinson’s disease^22^. SGTA plays crucial roles as a co-chaperone molecule, determining the fates of misfolded membrane and secretory proteins through a quality control mechanism regulated through interactions with the BAG6 complex^23,24^, and may influence early cellular events contributing to β-amyloid aggregation and toxicity^25^.

Expanding beyond comparison with AD diagnosis, we computed similar correlations for all model factors in both species (**Fig. 2B**). As before, we included genes significantly altered in human iNs for each factor under study (*p* < 0.05) that were also measured in the marmoset fibroblasts. While no additional factors were correlated with *PSEN1* effects, we found that the sex effects from the marmoset fibroblasts were significantly correlated with neuropathology measures in human iNs. Human donor CERAD score (ρ = −0.474, *p* = 5.42 × 10^−7^ for 101 genes) and Braak stage (ρ = −0.177, *p* = 5.83 × 10^−3^ for 241 genes) were both inversely correlated with male signatures in marmoset fibroblasts, although this correlation was not present in the proteome.

### 3.3 Differentially expressed genes and proteins highlight disease-relevant functional processes

After multiple testing correction, 18 genes and 42 proteins were found to be differentially expressed in males, and 50 genes and 76 proteins were similarly altered in fibroblasts from *PSEN1* mutant marmoset donors. Of these, four genes and five proteins showed significant changes in expression levels (FDR < 0.1) both based on sex and *PSEN1* genotypes. The differentially expressed genes included *GLOD4* and *LAMB2*, which were overexpressed in wild-type males, as well as *PTPN5* and *CD74*, with higher levels catalogued for *PSEN1* females (**Fig. 3A**). The distribution for differentially expressed proteins (S100A10, TMOD3, GPX4, COL12A1, and PCDHGA6) was comparatively more uniform, with wild-type males generally showing the highest expression, and *PSEN1* females showing the lowest (**Fig. 3B**).

**Figure 3:**
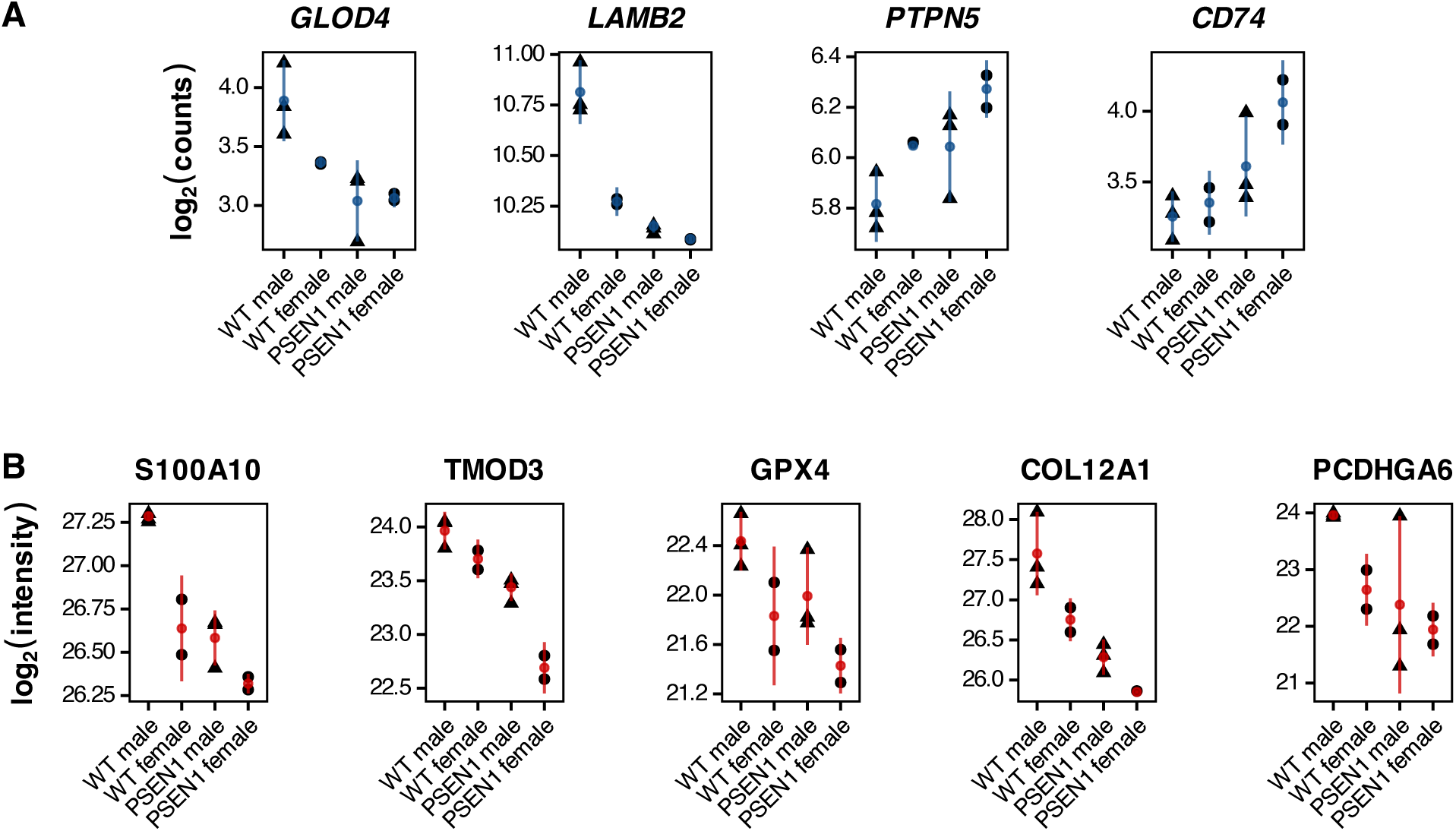
Genes and proteins differentially expressed both based on sex and *PSEN1* genotype. **(A)** Differentially expressed (FDR < 0.1) genes are shown in blue, with **(B)** proteins shown in red per genotype and sex group. Error bars show mean expression and 95% confidence intervals.

While these genes and proteins are not previously annotated to common biological processes, they are implicated in interconnected pathways of disease. Two genes, namely *GLOD4* and *LAMB2*, are altered in AD and have been noted to interact with toxic protein aggregates contributing to AD pathophysiology^26–28^, whereas others like *CD74* have been observed to be upregulated in activated microglia associated with AD and related cognitive decline^29–31^. *PTPN5* has been shown to mediate synaptic plasticity, showing increased levels in AD due to aggregating soluble β-amyloid oligomers^32^, much like what is observed in fibroblasts from *PSEN1* marmoset donors. COL12A1, a key component of the extracellular matrix (ECM) and found in fibroblasts in the brain, has decreased levels in cerebrospinal fluid (CSF), significantly correlating with pathological measures of AD^33^. GPX4 plays an important role in redox homeostasis by preventing lipid peroxidation, both of which are known to be impacted in Alzheimer’s disease^34^.

### 3.4 Transcriptomic alignment of marmoset fibroblasts and postmortem human co-expression modules reveals AD-relevant immune signature

To determine how the transcriptomic effects of *PSEN1* mutations in marmosets relate to human AD, we correlated RNA expression changes in marmoset fibroblasts to differences in human AD cases and controls across genes within a set of 30 co-expression modules derived from postmortem brain tissues (**Fig. 4A**). These 30 Alzheimer’s associated modules were derived from three cohorts and seven brain regions, and were then categorized into five consensus clusters based on common module gene content, function, and cell types^12^. The meta-analysis modules serve as a postmortem brain reference of gene expression changes in Alzheimer’s disease. We systematically compared the effects of sex and *PSEN1* mutations in marmoset fibroblasts to the effects of an Alzheimer’s disease diagnosis in the AMP-AD data by testing the correlation across common genes within each module.

**Figure 4:**
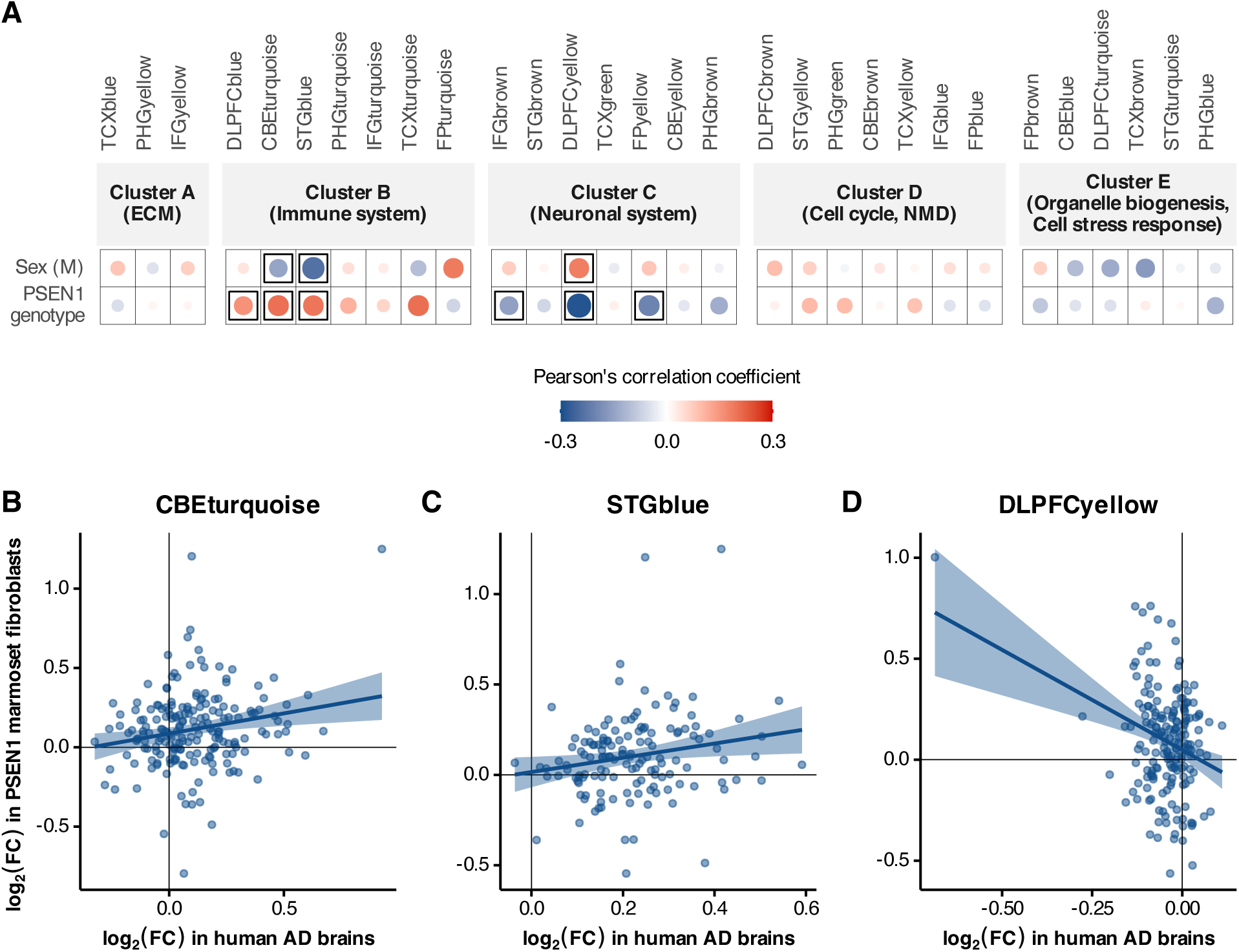
Transcriptomic changes highlight immune activation in fibroblasts derived from *PSEN1* mutant marmosets. **(A)** Correlations between gene expression effects of male sex and *PSEN1* genotype in the marmoset fibroblasts with AD case-control differences in 30 transcriptomic co-expression modules derived from post-mortem human brains. Disc size and intensity indicate the strength of the correlation, and significant (*p* < 0.05) correlations are framed. **(B-D)** The three modules with strongest correlations between human AD (x-axis) and marmoset *PSEN1* genotype (y-axis). Each dot corresponds to a resident module gene.

We found that the effects of a *PSEN1* mutation in marmoset fibroblasts were positively associated with human Alzheimer’s effects in Consensus Cluster B, which represents the immune system. Specifically, there were significant correlations with the modules DLPFCblue (177 genes, ρ = 0.155, *p* = 0.04), CBEturquoise (**Fig. 4B**; 193 genes, ρ = 0.202, *p* = 0.005), and STGblue (**Fig. 4C**; 141 genes, ρ = 0.193, *p* = 0.022). These modules represent similar immune signatures from three distinct AMP-AD study cohorts. We also found significant negative correlations for two of these modules, CBEturquoise (ρ = −0.146, *p* = 0.043), and STGblue (ρ = −0.235, *p* = 0.005), with male sex of the marmoset donor. While the limited number of genes on the NanoString panel precluded enrichment for key biological pathways or processes, the modules within Consensus Cluster B are typically associated with cytokine signaling and extracellular matrix interactions^8^. The DLPFCyellow module, belonging to Consensus Cluster C indicating the neuronal system, exhibited a significant negative correlation with human AD effects for *PSEN1* mutations (**Fig. 4D**; 185 genes, ρ = −0.286, *p* = 7.8 × 10^−5^) and a positive correlation with male sex (ρ = 0.182, *p* = 0.013) in marmoset fibroblasts.

The broader findings outlined through this analysis are consistent with previous studies highlighting differential module involvement at early (4 months) and later (12 months) ages in mouse models of AD^4^. Specifically, we note that the modules CBEturquoise, STGblue, and DLPFCyellow are found to be significant in both older male mice and in male marmoset donor fibroblasts with the same direction of correlation as shown here (**Fig. 4A**). Immune modules significant for *PSEN1* genotype, namely CBEturquoise and STGblue, were similarly positively correlated with 5xFAD mice, which are a transgenic amyloid model commonly used for studying AD in the laboratory. Module-level trends also highlight larger expression changes for genes in marmoset fibroblasts that follow the same trajectory as humans, like *SRGN* in CBEturquoise (**Fig. 4B**; top right in quadrant 1, log_2_ fold change in *PSEN1* marmoset fibroblasts = 1.25, and in AD human brains = 0.926), as well as those that differ in direction, namely *VGF* in DLPFCyellow (**Fig. 4D**; top left in quadrant 2, log_2_ fold change in *PSEN1* marmoset fibroblasts = 1.003, and in AD human brains = −0.686). Serglycin or *SRGN* is an intracellular heparan sulfate proteoglycan, found to be upregulated in brains with moderate to severe AD^35^. *SRGN* has also been identified as a disease-associated microglial gene, overexpressed in late AD^36^, likely driving microglial-dependent neuroinflammation^37^. *VGF*, a nerve growth factor (NGF) inducible gene, has been shown to be downregulated in AD in association with cognitive decline and neuropathology^38,39^, and multiscale causal network modeling approaches have identified it as a key driver in AD^40^. Altogether, our results indicate that fibroblast cultures *in vitro* can capture similar sex- and disease-specific changes as those observed in *in vivo* disease model systems, whilst calling attention to differing mechanisms underlying potential drivers of disease.

### 3.5 Proteomic alignment of marmoset fibroblasts and postmortem human co-expression modules reveals alterations in cellular metabolism

As with RNA expression, we systematically compared marmoset fibroblast proteomic signatures to 44 protein co-expression modules derived from postmortem human brains, 16 of which were associated with AD diagnosis and neuropathology^13^. As with the transcriptomic co-expression modules above, we assessed correlations between the effects of male sex and *PSEN1* mutations in marmoset fibroblasts and changes in human cases versus controls across the proteins in each module common to both data sets (**Fig. 5A**).

**Figure 5:**
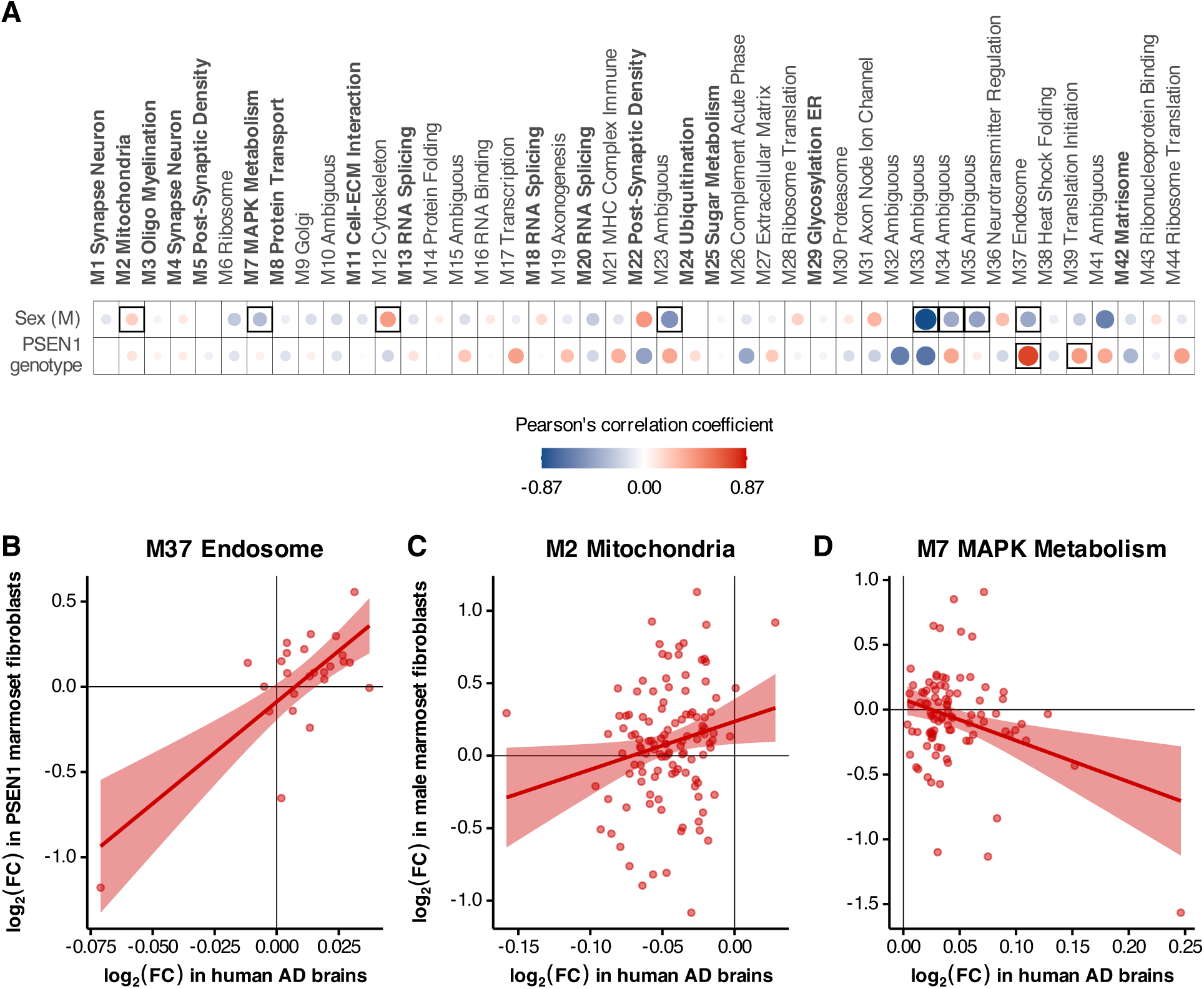
Marmoset fibroblast proteomes indicate changes in basic cellular processes and signaling. **(A)** Correlations between protein expression effects of male sex and *PSEN1* genotype in the marmoset fibroblasts with AD case-control differences in 44 proteomic co-expression modules derived from post-mortem human brains. The 12 modules labeled in boldface were associated with AD diagnosis and/or cognitive decline. Disc size and intensity indicate the strength of the correlation, and significant (*p* < 0.05) correlations are framed. **(B)** Protein expression effects for the Endosome module (M37) from *PSEN1* mutation in marmosets and the (**C**) Mitochondria (M2) and **(D)** MAPK Metabolism (M7) modules for male sex in marmoset donors (y-axis) versus human AD case-control differences (log_2_ fold change; x-axis). Each dot corresponds to a resident module gene.

We did not observe statistically significant correlations with any of the 16 AD modules for *PSEN1* genotype. However, across all 44 modules, *PSEN1* effects did correlate with postmortem AD in the endosome (M37) (**Fig. 5B**; 25 proteins, ρ = 0.74, *p* = 2.35 × 10^−5^) and translation initiation (M39) modules (31 proteins, ρ = 0.41, *p* = 0.021). Although the endosome module did not correlate with cognitive decline in the human study, it was positively associated with amyloid, tau, and TDP-43 neuropathology^13^. This correlation therefore suggests that the potentially amyloidogenic *PSEN1* effects in marmoset fibroblasts may recapitulate neuropathological signatures in the endosomal proteome.

Male marmoset sex effects were significantly associated with AD in multiple modules, including negative correlations with the M7 MAPK metabolism module (**Fig. 5D**; 91 proteins, ρ = −0.31, *p* = 0.0027), and several modules with ambiguous functional roles (M23, M33, M34, M35, with ρ < −0.39 and *p* < 0.05). Additionally, a positive association was found with the M2 Mitochondria (**Fig. 5C**; 117 proteins, ρ = 0.2, *p* = 0.029) and M12 cytoskeleton modules (33 proteins, ρ = 0.41, *p* = 0.018). The endosome module (M37) was also found to negatively correlate with male effects (ρ = −0.41, *p* = 0.043), showing distinct effects from *PSEN1* mutations (**Fig. 5A**). Both the M2 Mitochondrial and M7 MAPK metabolism modules are strongly enriched for proteins found to colocalize with neurofibrillary tangles and β-amyloid plaques, and along with some ambiguous modules like M23 and M33, are known to mediate cognitive outcomes^13,41^. Furthermore, many modules achieving statistical significance in our analysis were those that were not found to be preserved in transcriptomic networks^13^, namely modules M7, M34, M35, M37, and M39; highlighting perturbations detectable solely at the proteomic level.

Analyzing module-level relationships helps highlight the extent of changes in protein expression between the marmoset and human system. For instance, SDHD in the M37 Endosome (**Fig. 5B**; bottom left in quadrant 3, log_2_ fold change in *PSEN1* marmoset fibroblasts = −1.18, and in AD human brains = −0.07), and SLC38A2 in the M7 MAPK Metabolism modules (**Fig. 5D**; bottom right in quadrant 4, log_2_ fold change in male marmoset fibroblasts = −1.57, and in AD human brains = 0.25), show relatively larger fold changes in marmosets as compared to humans. SDHD or succinate dehydrogenase subunit D forms a part of mitochondrial Complex II, and is involved in both the TCA cycle and the electron transport chain, thus playing a crucial role in aerobic cellular respiration for energy production. Our results are consistent with numerous studies noting that subunits of mitochondrial complexes, including SDHD, are downregulated in both early-onset^42,43^ and late-onset Alzheimer’s disease^44^, as well as in amyloid mouse models of AD like the humanized Aβ knock-in^45^ and 5xFAD-Dp16, a Down syndrome-associated AD model^46^. SLC38A2, on the other hand, is a sodium-coupled amino acid transporter that allows cellular uptake of glutamine amongst others, needed for maintaining homeostasis in amino acid levels and for neurotransmitter synthesis. SLC38A2 has been implicated in selective hippocampal vulnerability in association with Alzheimer’s pathology and has been shown to be upregulated in AD brains^47,48^. However, our results are more in line with an observed downregulation in SLC38A2 levels in C57BL/6 mice injected with Aβ_1-42_, for an acute model of amyloid toxicity, highlighting it as a metabolic node supporting redox homeostasis; dysregulation of which leads to a disruption of the GSH-GPX4 antioxidant axis^49^. Indeed, in line with this idea, we also see reductions in GPX4 protein levels in *PSEN1* marmoset fibroblasts (**Fig. 3B**; β = −0.337, FDR = 0.092). We are thus able to detect key metabolic alterations at the protein level consistent with early-onset or amyloid-specific pathology in Alzheimer’s disease, even in non-neuronal contexts.

## 4. DISCUSSION

In this study, we profiled fibroblast cell lines derived from skin biopsies of marmosets targeted for genetic editing of *PSEN1* with sex- and age-matched wild-type controls. Since the donor animals were founders, the fibroblast lines contained a variety of *PSEN1* lesions. Although these cells were not reprogrammed and subsequently differentiated to cell types found in the brain, we nevertheless found relevant gene and protein expression changes common with those observed in human iPSC-derived neuronal lines derived from donors with AD pathology^11^. These findings suggest that *PSEN1* mutant marmoset fibroblasts capture part of the molecular effects typically seen in individuals diagnosed with sporadic or late-onset AD as represented in the ROSMAP cohort.

A number of fibroblast-expressed genes and proteins that surfaced in our analyses were of clear disease relevance. We do not observe any significant changes in gene or protein expression for APP, likely due to donor age and cell identity, however, many gene products identified to have significantly altered expression have been associated with AD pathology in literature. More specifically, we seem to converge onto distinct biological processes affected by the presence of amyloid in AD, across both RNA and protein modalities. Some differentially expressed genes like PTPN5 show increased expression due to activation by Aβ, which takes place even before plaque formation, and underlies early synaptic deficits that are commonly seen in mouse models of amyloidopathy^50,51^. COL12A1 shows reduced protein expression in brains and CSF from AD donors, whose abundance significantly correlates with Aβ_1-42_ and measures of tau pathology in AD CSF^33^. As the marmoset mutants contain an engineered early-onset mutation in *PSEN1* that affects APP processing^7^, hits in related functional aspects of AD pathology were not unexpected. Interestingly, we were able to identify genes with direct links to pathology measured in the central nervous system, an unanticipated finding from skin fibroblast cultures.

We also observed sex-specific effects in our study. In general, effects for males typically showed opposite direction of effect to *PSEN1* genotype (**Figs. 4** and **5**). This pattern corresponds to common additive changes in females and *PSEN1* mutants, leading to the greatest separation between wild-type males and mutant females (**Fig. 3**). *PSEN1* females thus seem to show the most extreme effects. Given that many of the same co-expression modules were significant for both sex and *PSEN1* mutations, we speculated that the observed genotype effects might be sex specific or otherwise interactive. In particular, multiple studies have observed a greater risk of Alzheimer’s in women and sex-specific gene expression signatures. However, including a sex-by-*PSEN1* interaction in our models failed to yield significant associations, possibly due to our very limited sample size. Nevertheless, the consistent sex and genotype effects in key genes and proteins motivates future studies of AD-relevant sex-specific differences in the population for both mutant and wild-type marmosets.

Distinct biological mechanisms seem to drive alterations observed in transcriptomes and proteomes. At each level of analysis, we observed that different genes and proteins contributed to the similarities between *PSEN1* genotype and human AD diagnosis. Specifically, although 32 genes and 45 proteins that were differentially expressed in the ROSMAP iNs for AD cases as compared to controls were measured in *PSEN1* marmoset fibroblasts, none of the differential proteins were products of the differential genes. This suggests that the *PSEN1* mutations engender distinct transcriptomic and proteomic signatures that drive the observed correlations between the mutations and AD status in humans. Furthermore, *PSEN1* marmoset fibroblasts mimicked RNA co-expression modules from human postmortem brains for neuronal and inflammatory processes, whereas the similar protein changes were in metabolism modules (**Figs. 4** and **5**). These findings are consistent with multi-omic results from human and mouse brains in which protein modules altered in AD are not present at the mRNA level^3,13^. That this disconnect is observed in undifferentiated fibroblasts further supports the growing consensus that transcriptomic and proteomic profiles often reveal fundamentally distinct and complementary aspects of disease, and reinforce the use of non-human primates as reliable research models.

Our study was limited by the number of genes and proteins that could be detected. In spite of only assaying a subset of all genes that can be measured through bulk RNA sequencing technologies, the NanoString AD panel served as a robust indicator of the perturbations in expression levels for genes associated with AD and related dementias (ADRD) for nonhuman primate models like the common marmoset; previously only validated for rodent models^8^.

Similarly, although our LFQ proteomics quantifies a fraction of proteins observed using tandem mass tag or data independent acquisition mass spectrometry, this sample was sufficient to yield disease relevant similarities. Surveying only fractions of the genome and proteome further reduced our ability to identify concerted changes in groups of genes or biological pathways; future studies will leverage genome- and proteome-wide technologies.

This pilot study motivates further disease modeling using cells derived from mutant marmosets. Despite studying skin fibroblasts sampled from very young founder animals (all < 9 months), we were able to identify relevant links to disease biology that primarily mimicked the direction of changes reported in humans. We are currently propagating later generations of heterozygous *PSEN1* C410Y marmosets, which will be integral in supporting future studies to identify disease mechanisms. Additionally, in spite of comparing skin fibroblasts in marmosets with terminally differentiated cells originating from postmortem brains, where effect sizes tend to be larger, we were able to identify significant changes in AD-relevant biological processes. As even wild-type marmosets often show aging-associated and AD-like pathophysiology^5,6^, studying mutant and wild-type marmosets provides a means to pair full lifespan assessment with a tractable cellular modeling platform for *in vitro* validation. Potential avenues of investigation include reprogrammed iPSC lines, direct differentiation into brain-relevant lineages to conserve cell age, and advanced co-cultures that enable organoid modeling. Noting that the human co-expression modules linked to marmoset *PSEN1* fibroblast signatures were strongly associated with changes in global cognitive function, we are encouraged that these models may reveal the cellular and molecular processes that underlie age-related decline in the aging marmoset brain.

## ACKNOWLEGEMENTS

This study was supported by the National Institutes of Health grant U19AG074866. The authors are grateful to Dr. Julia Oluoch, Brianne Stein, Erica Griffith, and the team of dedicated marmoset veterinary and husbandry colleagues from the University of Pittsburgh Systems Neuroscience Animal Research Laboratory who provide exceptional care of the marmosets and assist with these studies.

The results published here are in whole or in part based on data obtained from the AD Knowledge Portal (https://adknowledgeportal.org). Data generation was supported by the following NIH grants: P30AG10161, P30AG72975, R01AG15819, R01AG17917, R01AG036836, U01AG46152, U01AG61356, U01AG046139, P50AG016574, R01AG032990, U01AG046139, R01AG018023, U01AG006576, U01AG006786, R01AG025711, R01AG017216, R01AG003949, R01NS080820, U24NS072026, P30AG19610, U01AG046170, RF1AG057440, and U24AG061340, and the Cure PSP, Mayo and Michael J Fox foundations, Arizona Department of Health Services and the Arizona Biomedical Research Commission. We thank the participants of the Religious Order Study and Memory and Aging projects for the generous donation, the Sun Health Research Institute Brain and Body Donation Program, the Mayo Clinic Brain Bank, and the Mount Sinai/JJ Peters VA Medical Center NIH Brain and Tissue Repository. Data and analysis contributing investigators include Nilüfer Ertekin-Taner, Steven Younkin (Mayo Clinic, Jacksonville, FL), Todd Golde (University of Florida), Nathan Price (Institute for Systems Biology), David Bennett, Christopher Gaiteri (Rush University), Philip De Jager (Columbia University), Bin Zhang, Eric Schadt, Michelle Ehrlich, Vahram Haroutunian, Sam Gandy (Icahn School of Medicine at Mount Sinai), Koichi Iijima (National Center for Geriatrics and Gerontology, Japan), Scott Noggle (New York Stem Cell Foundation), and Lara Mangravite (Sage Bionetworks).

## CONFLICTS OF INTEREST STATEMENT

S.K., A.H., C.S., D.D., L.B., S.H.C., S.H., Y.M., S.K.H., J.E.P., G.E.H., and A.C.S. report no competing interests to declare at the time of submission. G.W.C. has served as a consultant for Astex Pharmaceuticals. N.T.S. is a co-founder and board member of Emtherapro Inc.

## CONSENT STATEMENT

No human participants were included in the study.

